# Meat enriched-diet and inflammation promote PI3Kα-dependent pancreatic cell plasticity that limit tissue regeneration

**DOI:** 10.1101/2023.08.22.554245

**Authors:** C Cayron, D Bozoglou, A V Villard, G Reyes-Castellanos, N Therville, R Baer, S Arcucci, N Naud, M Tosolini, F Pont, D Ferreira Da Mota, F Guéraud, C Basset, A Carrier, F Pierre, B Thibault, M Dufresne, J Guillermet-Guibert

## Abstract

**Objective:** Increased consumption of meat is an epidemiologically validated risk condition for pancreatic cancer development, but the underlying mechanisms and whether it is related to induction of epithelial cell plasticity are unknown.

**Design:** Experimental protocol to test the influence of high consumption of meat was compared to pancreatic inflammation experimental models. To determine the molecular drivers promoting pancreatic cell plasticity, we compared transcriptomics data sets from human samples of pancreatic inflammation and pancreatic cancer (PDAC) prone to plasticity and validated in vivo, ex vivo and in vitro the main identified target.

**Results:** Meat-enriched diet promoted plasticity of pancreatic acinar cells, that transdifferentiated in duct-like cells, and presented PI3K activation. We identified a selective PI3K activation gene signature enriched with plasticity. In this signature, *PHGDH*, which encodes an enzyme responsible for amino acid serine synthesis, was differentially expressed. High level of PHGDH in acinar cells was necessary for the proliferative action of PI3Kα sustained by an increased maximal mitochondrial capacity and decreased cyclin-dependent inhibitor p27 level. PHGDH level was decreased in transdifferentiated acinar cells. In this context, active PI3Kα promoted cell plasticity but decreased the number of cycling cells. Both epithelial-restricted genetic inactivation of PI3Kα and full PI3Kα inhibition by pharmacological dosage reduced inflammation-induced tissue damage, while a pharmacological PI3Kα activator promoted PanIN precancer lesion development.

**Conclusion:** Meat-enriched diet promoted plasticity. Blockage of plasticity by PI3Kα inhibition provoked an increased rate of acinar cell proliferation that had a beneficial impact on the tissue microenvironment less prone to precancer lesion development.

**What is already known on this topic:** It is now well accepted that inflammatory conditions predispose to pancreatic tumour development; increased consumption of red and processed meat is an epidemiologically validated risk condition, but the underlying mechanisms are unknown.

**What this study adds:** We identify PI3K activation as a common molecular pathway activated by increased consumption of red and processed meat and by inflammatory condition to promote pancreatic plasticity and precancer lesion development.

**How this study might affect research, practice or policy:** As we show that treatments with the clinically available PI3Kα inhibitor block pancreatic plasticity under inflammatory stress while maintaining pancreas mass and limiting inflammatory reaction damage, they may represent an efficient and safe preventive interception drug in patients at risk of developing pancreatic cancer. PI3K pro-cancer action is exacerbated by the loss of serine synthesis enzyme; hence, diets that alter amino acid synthesis should be tightly controlled in those patients.

## Introduction

Inflammatory conditions are generally thought to set the soil for cancer initiation and to share common molecular induction pathways with neoplasia, explaining the increased prevalence of cancers in patients with chronic inflammation. For patients with pancreatic inflammation [acute pancreatitis (AP), hereditary pancreatitis or chronic pancreatitis (CP) from all etiologies (alcoholic, hereditary, obstructive, autoimmune)], pancreatic cancer represents a serious complication (*1–3*). The burden of all these pancreatic disorders is expected to increase over time. Inflammation was shown to prevent oncogene-induced senescence and thus to be permissive for pancreatic cancer initiation (*4*).

During inflammation and initiation of cancer, common cellular events occur, such as the transdifferentiation of acinar exocrine cells into duct cells defining acinar-to-ductal metaplasia (ADM). Recently, it was demonstrated that the acinar-to-ductal plasticity promotes an epigenetic reprogramming, called epithelial memory of inflammation, which later favours cancer initiation (*5*). Pancreas is thus described as a plastic organ, where differentiated cells are facultative progenitors able to transdifferentiate in other differentiated cell types (*6–9*). Interestingly, not all acinar cells transdifferentiate in response to stress. The exocrine lineage also responds through increased proliferation (*6*). Unlike tissues that constantly renew themselves such as intestinal crypts, it is generally considered that no dedicated adult stem cells or stem cell niche are present in the pancreas to recover the tissue homeostasis and repair the damaged tissue. Instead, the proliferation of acinar cells that maintain their differentiation status helps to replace the damaged apoptotic acinar cells.

Understanding early molecular events of exocrine pancreas plasticity/proliferation balance occurring during pancreatic inflammatory conditions is critical to predict and support the treatment of patients at higher risk of developing pancreatic cancer. This topic is particularly important because pancreatic ductal adenocarcinoma (PDAC) is one of the top five causes of death by cancer and its incidence is on the rise, particularly in <65y population (*10–13*). It is now well accepted that there are also other inflammatory conditions than CP and AP that predispose to pancreatic tumour development, such as diabetes (*14–19*). Increased consumption of red and processed meat is an epidemiologically validated risk condition (*20*, *21*), but the underlying mechanisms and whether it is related to increased pancreatic inflammation or induction of ADM are unknown.

In this study, we identified that meat consumption promotes pancreatic plasticity and that blockage of plasticity by the mean of phosphoinositide-3-kinase PI3Kα inactivation promotes tissue regeneration by proliferation. We found that the clinically available PI3Kα inhibitors are compatible with their use as cancer prevention drugs in pancreatic inflammation.

## Results

### Meat consumption promotes tissue plasticity associated with PI3K activation and with an increase of quiescence markers

In an aim to uncover common grounds to various factors known to increase the risk of developing PDAC, we compared pancreas from rats fed with control diet or with a diet enriched in meat with a high proportion of red meat, known to increase the risk of PDAC (*20*). This later nutritive regimen is called from there Meat diet (Fig. 1a) and was associated selectively with the presence of ADM lesions in rat pancreas (Fig. 1b). Indeed, pancreatic ADM lesions indicative of increased acinar cell plasticity were observed only in the cohort of rats that had Meat diet (Fig. 1b). PI3Kα is critical for ADM formation and for the induction of pancreatic cancer by oncogenic KRAS in the presence of mutated p53 and / or concomitant inflammation (*22–24*). Meat diet increased the level of pS473AKT (Fig. 1c), indicative of PI3K activation, and also of αSMA, indicative of stromal cell activation (Fig. 1c). To analyse possible regeneration process by proliferation, the cyclin-dependent kinase (cdk) inhibitor p27 (*25*) level was assessed and, despite the increase of PI3K activation usually associated with decreased cdk inhibitor expression (*26*), p27 level was significantly increased (Fig. 1c).

**Figure 1:**
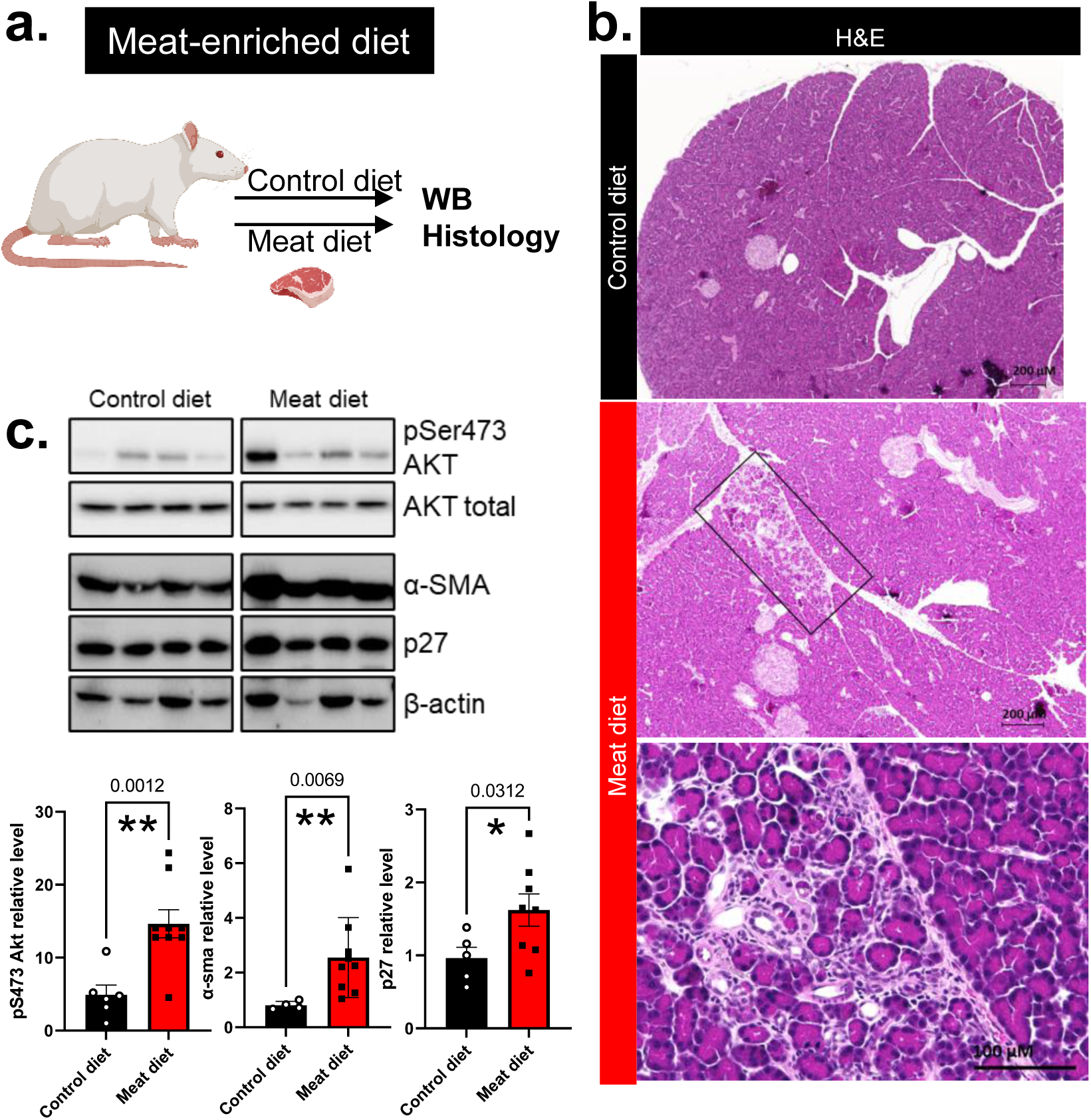
Meat consumption promotes tissue plasticity associated with PI3K activation and with an increase of quiescence markers. a. Experimental setup describing experimental diet enriched in red meat in rats. b. Representative H&E. c. WB and quantification of indicated proteins (n>3 in each group). pSerAKT levels are normalized with levels of AKT; other markers with β-actin. N≥3 biological replicates; Values are mean ± SEM. ANOVA tests: *:p<0.05; **: p<0.01; ***:p<0.001

These data suggest that pancreatic stress promotes both plasticity and regulation of cell quiescence marker that might favour tissue damage. Increased quiescence marker is maybe controlled by PI3K activation.

### PI3K/AKT pathway gene signatures and *PHGDH* are differentially enriched in chronic pancreatitis compared to pancreatic adenocarcinoma

To understand this observation and validate its pertinence in human disease, we sought to determine in an unbiased manner which molecular events were selectively associated with tissue-response to stress including acinar cell plasticity and cell proliferation, by comparing pancreatic inflammation and cancer. We analysed publicly available transcriptomic datasets to distinguish chronic pancreatitis (CP) from primary tumours of PDAC (PDAC) (Fig. 2a-c, Suppl. Table 1). CP and PDAC patients presented differential enrichment of 22 mRNA expression-based hallmarks of biological pathways, also enriched compared to normal pancreas. The PI3K/AKT/mTOR signalling pathway was found in the top five of the most differentially enriched with the lowest p-value (Fig. 2a). Differential enrichment analysis demonstrated that selective PI3K/AKT/mTOR pathway signatures were significantly enriched in CP and in PDAC compared to normal parenchyma (Fig. 2b): this is what emerged from Reactome pathway analysis. In detail, we found that PI3K cascade to FGFR1-4 were significantly increased in CP as opposed to PDAC and normal parenchyma, suggesting differential activation of receptor tyrosine kinase (RTK)-coupled PI3K in these samples (Fig. 2b). Reversely, PI3K cascade to ERBB2, that has known role in PDAC initiation downstream KRAS (*27–31*), was significantly increased in PDAC as opposed to CP and normal parenchyma.

**Figure 2:**
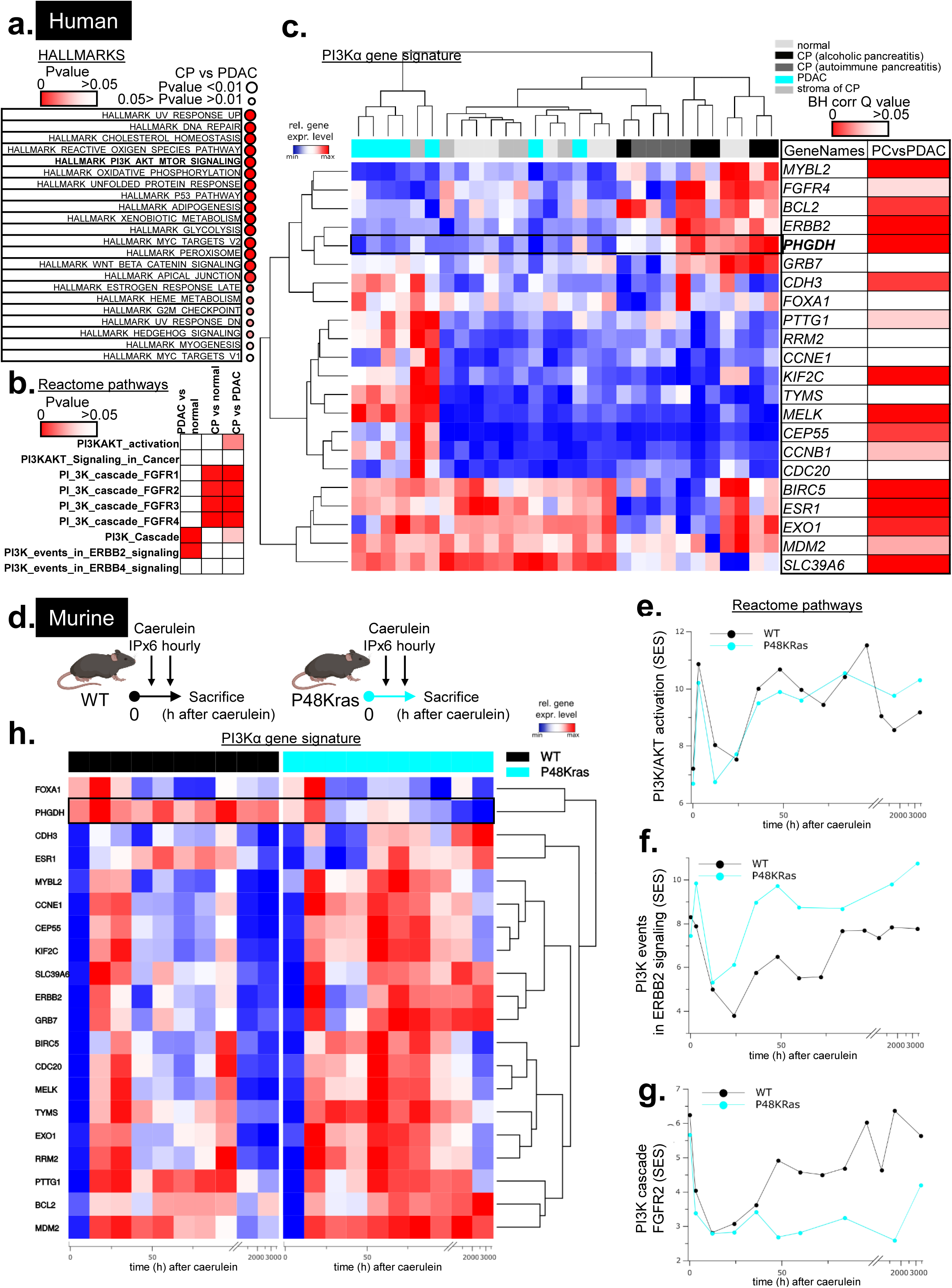
PI3K/Akt pathway and PHGDH are differentially enriched in chronic pancreatitis compared to pancreatic adenocarcinoma. **a.-c.** We used raw data from various transcriptomic studies (*58–60*) with compatible platforms and re-analysed them. **a.** An unbiased search of HALLMARKS differentially expressed in CP vs PDAC patients was performed; only hallmarks with p value <0.05 are shown. **b.** Statistical analysis (p values of enrichment scores) of Reactome PI3K pathway enrichment in PDAC vs normal, CP vs normal or CP vs PDAC is shown. Intensity is shown with a threshold at 0.05. **c.** Clustering of human PI3Kα gene expression signature was performed by R in normal, CP (alcoholic pancreatitis, autoimmune pancreatitis), pancreatic ductal adenocarcinoma (PDAC) and microdissected stroma from chronic pancreatitis of undocumented aetiology. Blue = low expression, Red = high expression. The unsupervised hierarchy of each sample is shown in the top part of the figure; the unsupervised hierarchy of genes is shown on the left; the list of genes is detailed on the right, with statistical differences between PDAC and CP samples. BH-corrected Q values are represented. Statistical analysis is detailed in Supplementary Table 1. **d.-h.** We used the raw data of a time course evolution of gene expression previously performed in WT (black line) or KrasG12D (blue line) genetic contexts upon caerulein-induced inflammation (*34*) and reanalysed gene expression in time. Times 0.5h-3h corresponds to early genes, 6h-48h middle genes, 48h-340h late genes. **d.** Scheme. **e.-g.** An unbiased pathway analysis (Reactome) was performed and, for each reactome lists presenting PI3K in their title, SES enrichment scores are represented in time. Doted rectangle points out the time points at which the differential signature is observed between the two conditions. **h.** A heat map of gene expression (PI3Kα signature) shows the differential expression of these genes in time in the two genetic contexts. *<u>PHGDH</u>* is high only in WT background upon Caerulein treatment (highlighted line), at a time point when FGFR1/4-PI3K signatures are significantly increased in this genetic context.

Considering that PI3Kα is key for RTK signalling (*32*) and PDAC initiation and progression (*22*, *24*), we then used the previously published PI3Kα activation gene signature, based on expression levels of PI3Kα activation-associated curated genes (*24*). PI3Kα activation scoring allowed us to cluster together all CP patients from two different aetiologies (Fig. 2c). Conversely, PDAC clustered mostly with activated stroma of CP. Amongst the list of genes in the signature, *PHGDH* was highly expressed in the CP-rich cluster. PHGDH (phosphoglycerate deshydrogenase) is an enzyme at the crosstalk between glucose and amino-acids metabolism. It oxidizes the glycolytic intermediate 3-phosphoglycerate (3-PG) into 3-phosphohydroxy pyruvate (3-PHP) (rate-limiting step in serine synthesis from 3-PG) and controls ROS detoxification (*33*).

To confirm the relevance of our experimental rodent models to study this process, we compared gene expression upon two sequential injections of caerulein (*34*), a stable form of cholecystokinin (CKK) that promotes pancreatic inflammation, in mice non harbouring (mimicking inflammatory condition, a condition at risk for developing cancer) or harbouring mutant KRAS^G12D^ (mimicking early steps of cancer) (Fig. 2d). The same evolution of gene signatures compared to human data set was observed. Indeed, while the generic PI3K/AKT activation signature was evolving in a similar manner in both groups of mice (Fig. 2e), the selective PI3K events in ERBB2 signalling or the PI3K events in FGFR2 signalling were significantly enriched in mutant KRAS (precancer) or in WT pancreas (inflammation only), respectively (Fig. 2f,g). As found in human CP, the expression of *PHGDH* was enriched differently in WT in comparison to KRAS-oncogenic model (Fig. 2h). After caerulein treatment, WT mice show a high and persistent PHGDH expression level, whereas in Krasmut mice the PHGDH level is transient and decreases in the long term.

Hence, both chronic pancreatitis and PDAC present differential PI3K/AKT and PI3Kα activation signatures. Amongst the genes which expression is associated with PI3Kα activity, *PHGDH* levels were increased in CP and inflammation only condition, suggesting a role in pancreatic stress. In addition, the experimental models that we used are pertinent in reproducing those molecular alterations observed in human.

### PHGDH protein level is decreased in plastic acinar cells

We next focused on PHGDH which was amongst the most differentially expressed molecular markers identified in Fig. 2, and analysed its protein level by Western blotting and histology. PHGDH protein level was increased by meat diet in rats (Fig. 3a). Interestingly, PHGDH immunohistochemistry analysis showed increased levels of PHGDH induced by caerulein in mice only in acinar cell that maintained their identity and not in ADM structures in pancreas (Fig. 3b,c).

**Figure 3:**
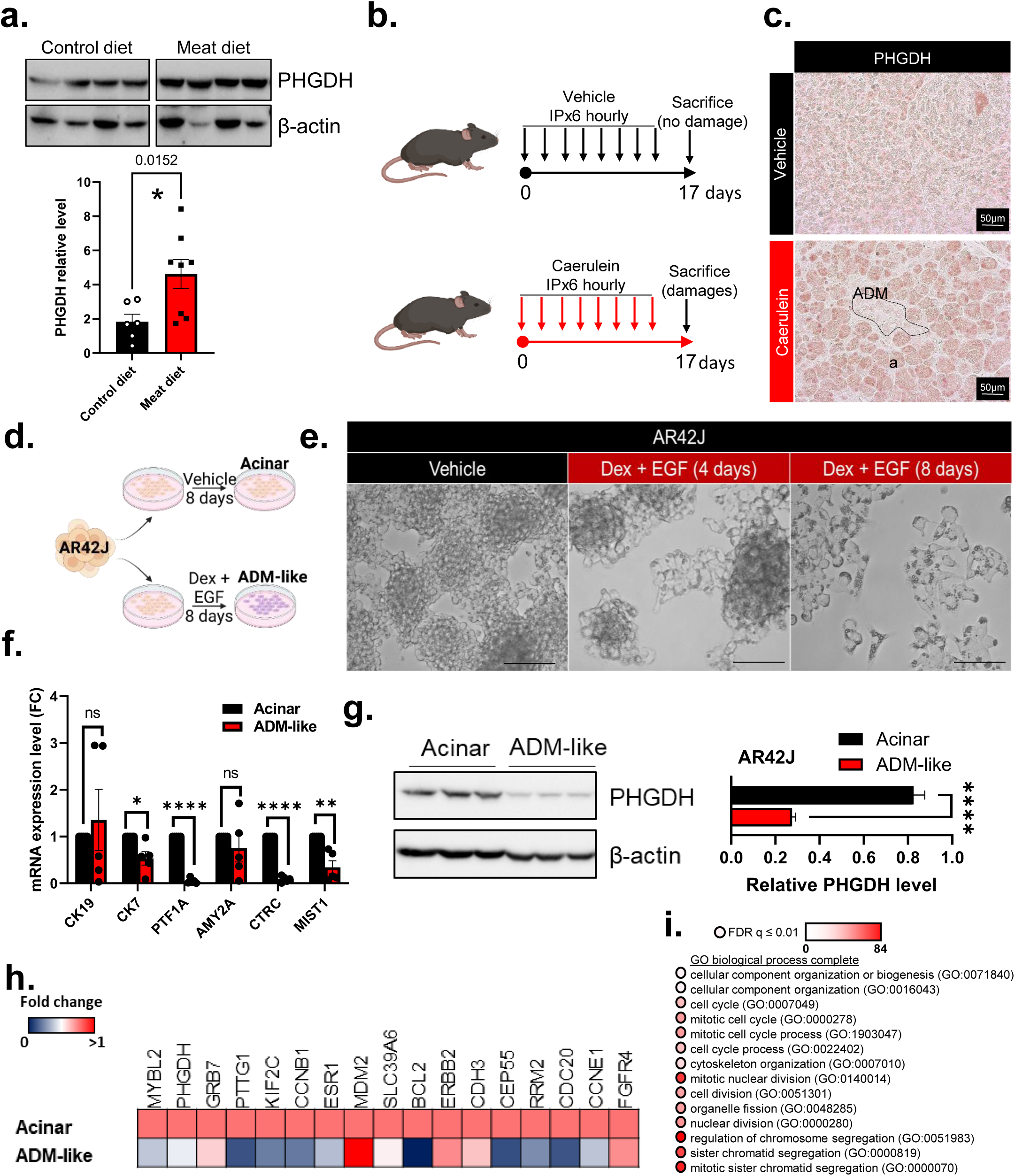
PHGDH protein level is decreased in plastic acinar cells. **a.** WB and quantification of indicated proteins (n>3 in each group) in pancreas from diet enriched in red meatcohorte. PHGDH level is normalized with β-actin. N≥3 biological replicates; Values are mean ± SEM. ANOVA tests: *:p<0.05; **: p<0.01; ***:p<0.001 **b.** Experimental setup describing experimental chronic pancreatitis model in mice. **c.** IHC against PHGDH in vehicle vs. caerulein-injected pancreas: a representative image is shown (n=3 in each group). a=acini; ADM=acinar-to-ductal metaplasia **d.** AR42J-B13 cells were treated or not (Vehicle named acinar) for 8 days with Dex + EGF (named ADM-like). **d.** Representative morphology is shown. **f.** mRNA levels of indicated target genes (duct-specific genes: *CK19, CK7* ; acinar-specific genes: *PTF1A, AMY2A, CTRC, MIST1*) were assessed by RT-qPCR in acinar vs ADM-like cells. (n=5 independent experiments). **g.** Protein levels of indicated proteins were assessed by WB in ADM-like relative to acinar cells; quantification is normalised with β-actin levels (n=3 independent experiments). **h.** mRNA levels of indicated target genes (PI3Kα activation signature) were assessed by RT-qPCR in acinar vs ADM-like cells. (n=5 independent experiments). **i.** Gene ontology PANTHER analysis of PI3Kα gene signature. Intensity False Discovery Rate is shown in scaled colors. N≥3 independent experiments; Values are mean ± SEM. T-Test compared to Vehicle (2 conditions) or ANOVA tests (multiple conditions): *:p<0.05; **: p<0.01; ***:p<0.001

To test the importance of PHGDH decreased level in PI3K-controlled pancreatic plasticity, we set up an *in vitro* model, using rat acinar AR-42J cells treated with a cocktail of EGF and dexamethasone (Dex + EGF) every 2 days that mimics the biochemical cues found *in vivo* in stressed pancreas and that promotes their plasticity from acinar to duct-like cells (*35*). After 8 days of treatment, morphology changes were associated with loss of acinar cell markers (*CK7*, *PTF1A*, *CTRC*, *MIST1*) without increased expression of ductal marker (*CK19*) (Fig. 3d-f), indicative of an intermediate stage of metaplasia, from now called ADM-like. PHGDH level was decreased in ADM-like cells (Fig. 3g) as observed *in vivo* (Fig. 3c), further validating the relevance of our experimental setup.

We also found that some mRNA levels of PI3Kα signature (*PTTG1, KIF2C, CCNB1, BCL2, CEP55, RRM2, CDC20, CCNE1)* were decreased in ADM-like cells (Fig. 3h); these genes corresponded globally to the genes decreased in PC patients (Fig. 2c), while the gene ontology analysis of this gene list revealed association with pathways promoting proliferation (e. g. cell cycle (GO:0007049); Fig. 3i). These data led us to the hypothesis that activation of PI3K and altered levels of PHGDH could determine the level of proliferative regeneration versus plasticity in exocrine cells in stress conditions.

### Plasticity-blocked acinar cells maintain their proliferative capacity by preventing PHGDH positive control of p27 quiescence marker

To test this hypothesis, treatment with PI3Kα inhibitor GDC-0326 decreased the level of pSer473AKT in both acinar and ADM-like cell context (Fig. 4a,b), suggesting that both cell states harbour a PI3Kα activity. The level of cdk inhibitor p27 (quiescence marker) was increased by GDC-0326 treatment in acinar cells (Fig. 4a) but not in ADM-like cells (Fig. 4b). Thus, the action of PI3Kα inhibition on p27 level was observed only in PHGDH-high acinar cells, and not in cells in their intermediate metaplasic status. Reversely, genetically reduced levels of PHGDH by transfection of pools of siRNA targeting *<u>PHGDH</u>* in acinar cells decreased p27 levels to levels found in ADM-like cells (Fig. 4c) and mimicked DEX+EGF treatment. Finally, treatment with the pharmacological inhibitor of PHGDH (CBR-5884) in acinar cells that most efficiently inhibited PHGDH activity *in vitro* (Suppl. Fig. 1a) prevented the effect of GDC-0326 inhibitor on p27 level (Fig. 4d), further demonstrating causality between PHGDH level and PI3K output signalling. Both PI3Kα and PHGDH control mitochondrial activity (*33*, *36*, *37*), and oxidative phosphorylation signature was found in the top six of the most differentially enriched signatures with the lowest p-value in human CP compared to PDAC (Fig. 2a). We analysed mitochondrial activity with Seahorse technology in AR42J cells with high (acinar) and low (ADM-like) PHGDH levels and measured the effects of GDC-0326, a selective PI3Kα inhibitor and CBR-5884, a selective PHGDH inhibitor, alone or in combination (Fig. 4e-j). In cells with high PHGDH level (acinar), the combined treatment using GDC-0326 and CBR-5884 reduced significantly the basal respiration and the mitochondrial maximal capacity compared to these inhibitors alone (Fig. 4e,g,i), while the combinative treatment had little or no effect in cells with low levels of PHGDH (Fig. 4f,h,j).

**Figure 4:**
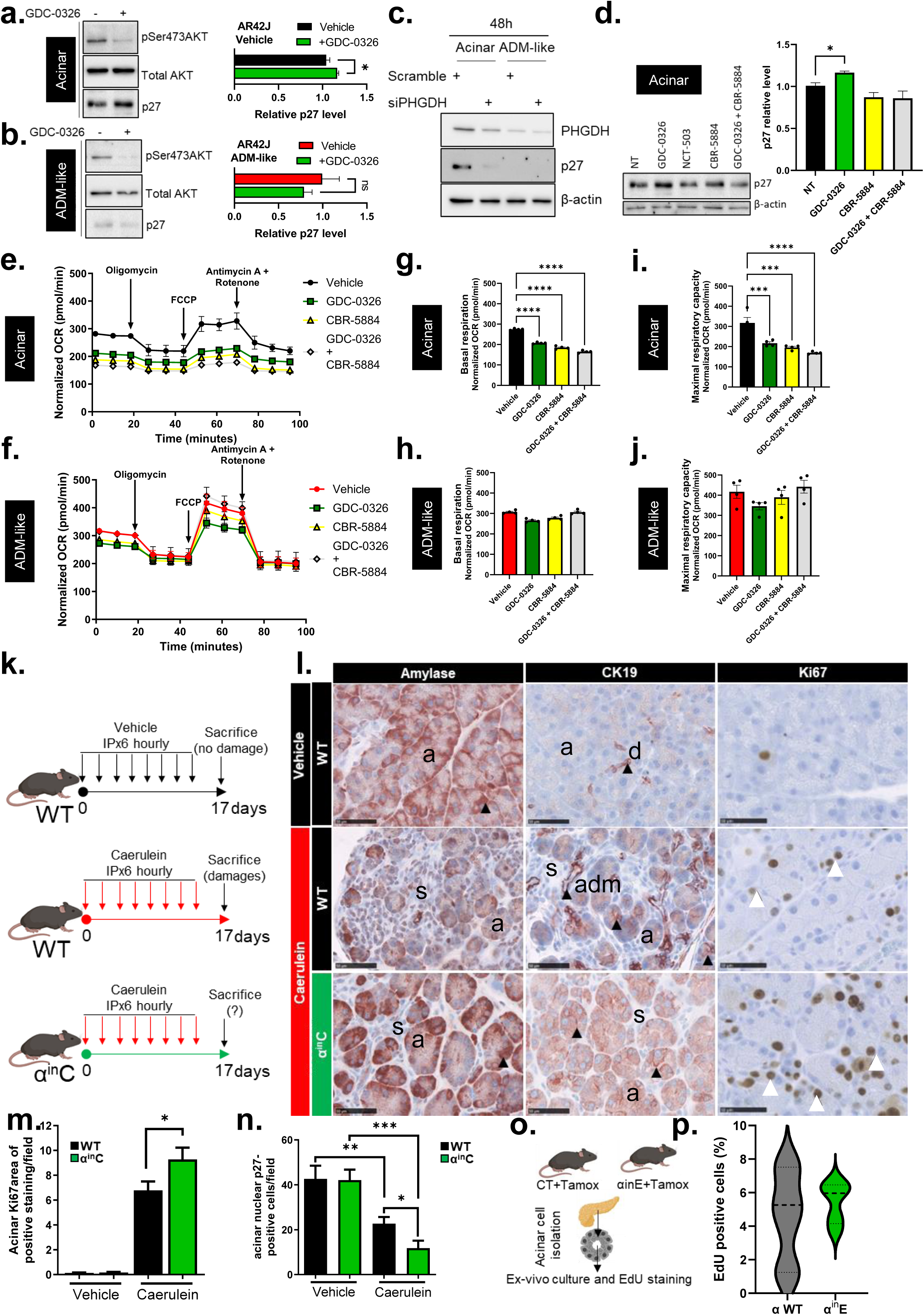
Plasticity-blocked acinar cells maintain their proliferative capacity by preventing PHGDH positive control of p27 quiescence marker. **a,b.** Protein and phospho-proteins levels of indicated target proteins were assessed by WB in acinar vs ADM-like cells; cells were treated or not with GDC-0326 (100nM) for 24h. (n=3). **c.** Protein levels of indicated target proteins were assessed by WB in Vehicle cells after transfection with scramble or PHGDH smart pools of siRNAs. **d.** Protein levels of indicated target proteins were assessed by WB in acinar cells after treatment with two PHGDH pharmacological inhibitors (NCT-503, 2.5µM; CBR-5884, 30µM). Cells were treated or not with GDC-0326 (100nM) for 24h. (n=3). **e-j.** Seahorse analysis indicative of glycolysis and mitochondrial activity levels was performed in 8-days Vehicle (acinar) vs Dex+EGF (ADM-like) cells. (n=4 independent experiments). Cells were treated or not with PI3Kα (GDC-0326, 100nM) or PHGDH inhibitor (CBR-5884, 30µM) for 24h. Data were normalized with the number of cells and between biological replicates. **k.** Experimental setup describing chronic pancreatitis model. **l-n.** IHC staining as indicated in WT or α^in^C pancreas subjected to the repeated injury chronic pancreatitis protocol (Caerulein). Scale: 20µm. Representative pictures are shown. a=acini; adm=acinar-to-ductal metaplasia; s=stroma; d=normal ducts. Black and white arrowheads indicate representative positive stainings for CK19 or Ki67, respectively. Positive cells or nuclei are quantified in five randomly pictured fields in each mouse. N≥4 mice. Values are mean ± SEM. T-Test: *=P<0.05. **o.** CT or α^in^E mice were injected with tamoxifen; 2.5 weeks after acini were prepared and analysed as indicated. Cre+ genotype was verified by PCR. **p.** After 2 days of culture, rate of EdU incorporation (2h pulse) was assessed by imaging with Operetta in CT+Tamox or α^in^E+Tamox acini with or without CBR-5884, 30µM. Quantifications are shown. (n>4). N≥3 independent experiments; Values are mean ± SEM. T-Test compared to Vehicle (2 conditions) or ANOVA tests (multiple conditions): *:p<0.05; **: p<0.01; ***:p<0.001

*In vivo*, we tested whether the sole genetic catalytic inactivation of PI3Kα altered the proliferation rate of ADM-like cells. This was possible by deleting the exons encoding PI3Kα catalytic domain (*37*, *38*) selectively in pancreatic epithelial lineage (called α^in^C model). Cre recombinase expression in Pdx1-positive cells induced the deletion of two exons encoding the catalytic domain of PI3Kα (Suppl. Fig. 1b). We subjected WT or α^in^C mice to eight series of caerulein injections over 17 days to induce a repetitive stress (Fig. 4k). Pancreatic cell-specific genetic inactivation of PI3Kα did not prevent caerulein-induced hypersecretion of acinar cells as measured by similar levels of plasmatic amylase at early time points (Suppl. Fig. 1c). PI3Kα inactivation led to decreased levels of pAKT substrate staining (Suppl. Fig. 1d). We analysed the caerulein effects on cells with acinar morphology. Caerulein-induced acinar cell atrophy was blocked by PI3Kα inactivation (Fig. 4l), and a moderate loss of amylase expression was observed in PI3Kα–deficient cells that maintained an acinar morphology compared to vehicle WT condition (Fig. 4l). The duct-specific marker cytokeratin 19 (CK19) was expressed in an increased number of duct-like structures in caerulein-treated WT cells but presented a moderate basolateral expression in cells with acinar morphology when PI3Kα was inactivated (Fig. 4l). This was associated with an increase of the proliferation Ki67 marker in caerulein-treated PI3Kα–deficient acinar cells compared to caerulein-treated WT acinar cells (Fig. 4k,m). We confirmed the action of PI3Kα inactivation on controlling quiescence marker p27 level and the effect were more pronounced *in vivo* than *in vitro* : p27 nuclear level was further significantly decreased by PI3Kα inactivation in caerulein-treated acinar cells (Fig. 4n, Suppl. Fig. 1e). These data show that under the stress promoted by repetitive injection of caerulein, PI3Kα catalytic inactivation largely maintains a proliferative regenerative response in the pancreatic exocrine parenchyma, which keep their acinar morphology.

We next used alternative approaches to confirm our results. We studied primary *ex vivo* cultures of acinar cells from murine pancreas, which are known to spontaneously transdifferentiate. Using ElastaseCreER+;p110αflox/flox mice (α^in^E), we genetically inactivated PI3Kα in adult acinar cells (*31*), promoting the recombination of two exons encoding p110α catalytic domain by CreER expression in Elastase-positive cells (acinar cells) and by tamoxifen treatment *in vivo*; control Cre negative mice (CT) were also tamoxifen treated. Exocrine cell preparation was performed, cultured *ex vivo* to promote spontaneous transdifferentiation, and EdU incorporation was analysed to monitor DNA replication assessing proliferation rates (Fig. 4o,p). The proliferation rate of ElastaseCreER-positive tamoxifen-treated pancreatic exocrine cells was homogenous between cell preparations and comprised only high proliferating acinar preparations compared to the ElastaseCreER-negative tamoxifen-treated pancreatic acini (Fig. 4p).

Taken together, these data demonstrate that accumulation of ADM-like cells upon PI3K inactivation maintains the proliferative capacity through a decrease in PHGDH level, suggesting that regeneration of the pancreas could be increased upon PI3Kα inactivation. PI3Kα inhibitors could block the cells in this cell state without prompting pancreas atrophy.

### Pharmacological or genetic down-regulation of PI3Kα activity in preclinical models of pancreatitis increases pancreas regeneration and reduces fibrosis

Finally, we aimed to test *in vivo* whether by blocking plasticity in PHGDH-low cells but not preventing their proliferation, PI3Kα catalytic inactivation could be beneficial to promote pancreas regeneration after inflammation. *In vivo*, a pharmacological inhibition of PI3Kα by GDC-0326 prevented the damage of repetitive injection of caerulein, while we only observed some isolated ADM and in very limited numbers (Fig. 5a,b). Interestingly, GDC-0326 treatment led to the absence of fibrosis accumulation (Fig. 5b, bottom, picrosirius staining). We next tested whether the sole genetic catalytic inactivation of PI3Kα in epithelial cells had an action on inflammatory damage intensity (Fig. 5 c-h). Epithelial-restricted PI3Kα inactivation decreased the number of ADM lesions (Fig. 5d). ADM occurs upon re-expression of key pancreatic progenitor factors, such as Sox9 (*7*). Nuclear Sox9 re-expression by caerulein was significantly decreased by genetic PI3Kα inactivation (Fig. 5e; Suppl. Fig. 2). Epithelial-restricted PI3Kα inactivation increased tissue damage protection in response to stress, quantified by a significantly decreased pancreatitis score (Fig. 5f) and fibrosis (Fig. 5g,h).

**Figure 5:**
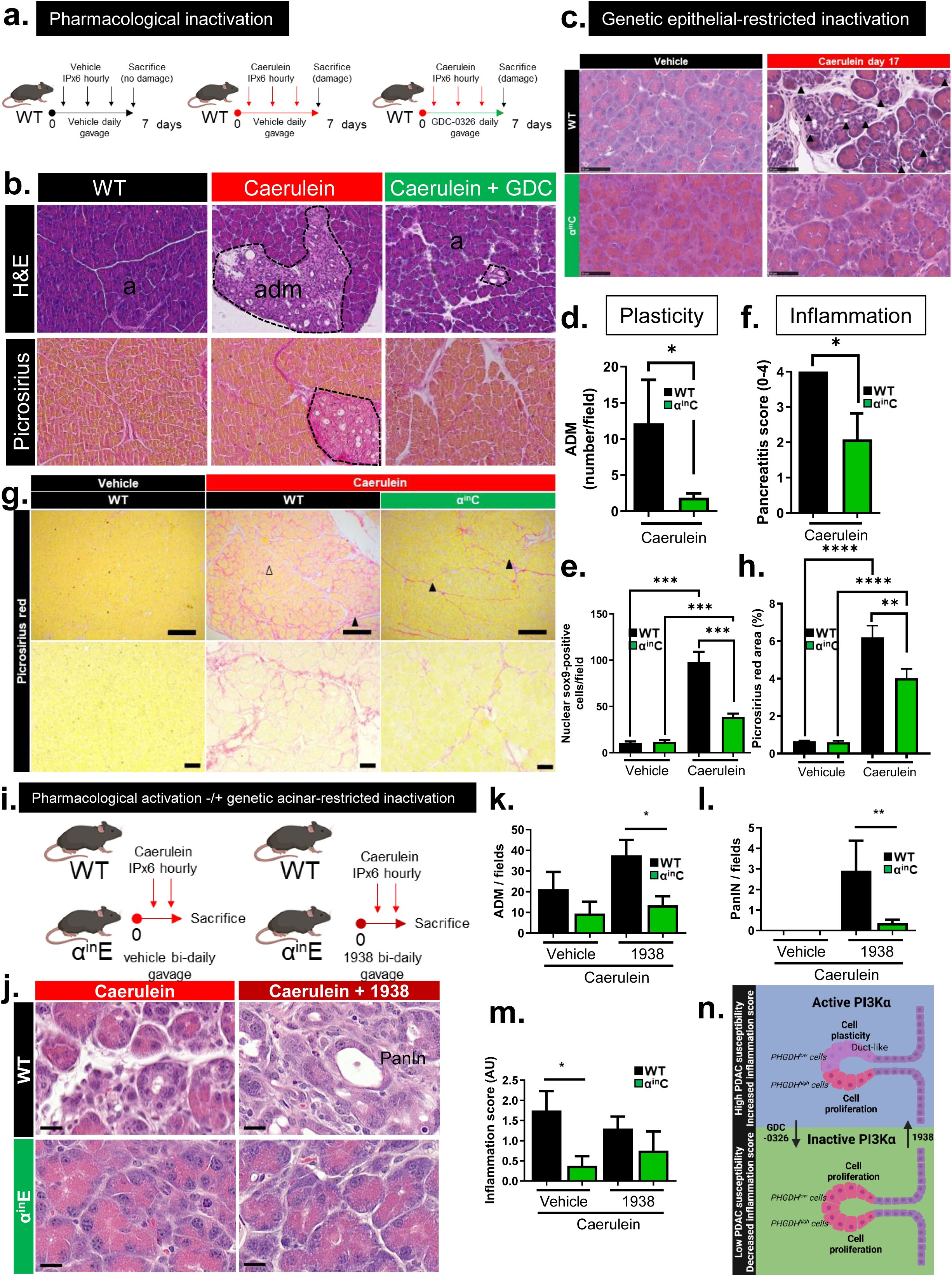
Pharmacological or genetic down-regulation of PI3Kα activity in preclinical models of pancreatitis increases pancreas regeneration and reduces fibrosis. **a,b.** Dosage schedule of PI3Kα-targeting drug (GDC-0326 10mg/kg daily-green arrow) in WT mice subjected to repeated pancreatitis protocol (red arrows) compared to vehicle treated mice (black arrows). n≥4 mice in each group. H&E and picrosirius coloration as indicated in vehicle- or GDC-0326-dosed and caerulein-treated WT pancreas. Representative pictures are shown. Scale bar=20µm. **c-h.** Scoring of ADM or pancreatitis score in Caerulein-treated pancreas harvested at day 17 from indicated genotype. This was performed with H&E stainings in WT or α^in^C pancreas subjected to the repeated injury chronic pancreatitis protocol (Caerulein). The proportion of pathological lobules / ADM was scored using 2 whole slide scans from each mouse, n=6 mice for each genotype. Values are mean ± SEM. T-Test: *=P<0.05. **g.** Picrosirius red staining as indicated in WT or α^in^C pancreas subjected to the repeated injury chronic pancreatitis protocol (Caerulein). Representative pictures are shown. Positive staining are quantified in five randomly pictured fields in each mouse. N≥4 mice. Values are mean ± SEM. T-Test: *=P<0.05. **e,h.** Indicated positive nuclei or area are quantified in five randomly pictured fields in each mouse. N≥4 mice. Values are mean ± SEM. T-Test: *=P<0.05. **i-m.** Dosage schedule of vehicle or PI3Kα-activating drug (1938 100mg/kg bi-daily black arrow) in WT+Tamox or α^in^E+Tamox mice subjected to two round of caerulein injection (red arrows). n≥4 mice in each group. H&E as indicated in vehicle- or 1938-dosed and caerulein-treated WT+Tamox or α^in^E+Tamox pancreas. Representative pictures are shown. Scale bar=20µM. Quantifications are shown as indicated. Values are mean ± SEM. T-Test: *=P<0.05; **=P<0.01. **l.** Graphical abstract.

We next decided to perform this experiment in a reverse way by increasing the level of PI3K activation. This was possible because, recently, a novel drug was developed to selectively activate PI3Kα activity (compound 1938, (*39*)). We dosed caerulein-treated mice with 1938 to test whether the response of the tissue to stress could be pushed towards increased cell plasticity by PI3Kα hyperactivation (Fig. 5i-m), and whether it increased pancreas damage as a consequence; to demonstrate the latter, we blocked ADM by genetically inactivating PI3Kα in pancreatic epithelial acinar cells (α^in^E genetic background). Acute treatment with 1938 promoted caerulein-induced metaplasic structures (Fig. 5j,k); genetic catalytic inactivation of PI3Kα restricted to acinar cells (α^in^E) prevented caerulein- and 1938-induced metaplasia (Fig. 5j,k). 1938 promoted the formation of PanIN lesions that were prevented by PI3Kα genetic inactivation (Fig. 5l). Finally, inflammatory score was unchanged by 1938 treatment in WT mice, while it was increased in α^in^E (Fig. 5j,m).

PI3Kα activity-dependent acinar cell plasticity, known to increase the risk to develop cancer, also drive fibrosis accumulation and inflammatory damage. Reversely, epithelial-restricted PI3Kα genetic or global pharmacological inactivation changes acinar cell fate under stress and promotes proliferative acinar cells that could also sustain pancreatic regeneration.

## Discussion

Determining the precise molecular links between chronic inflammatory disease, epidemiologically identified risk factor conditions and PDAC is necessary to predict which patients are at higher risk of developing this lethal disease. Here, we demonstrate that, in epithelial cells, the signalling enzyme PI3Kα positively contributes to a detrimental fate of acinar cells in repetitively injured pancreas by caerulein or by meat-enriched diet. PI3Kα activity both promotes acinar cell plasticity and prevents regeneration by cell proliferation (Fig. 5n, **Graphical abstract**). Mechanistically, PI3Kα activity and PHGDH levels cooperate in stressed acinar cells to control the cdk inhibitor p27 level and the fine tuning of mitochondrial activity.

Both PI3K and MAPK pathways are activated during pancreatic injury. PI3Kα controls actin-cytoskeleton remodelling necessary for ADM (*22*); while pancreas-restricted inhibition of MAPK pathway via short hairpins targeting MEK1/2 or their systemic inhibition via a pharmacological inhibitor trametinib (generic name) prevents both the formation of ADM and the turnover of pancreatic parenchyma, leading to a loss of pancreatic mass (*40*). It is interesting to note from our data that PI3Kα inhibition increases the turnover of pancreatic cells. The inhibition of PI3Kα signalling could potentially prevent the formation of precursor lesions in the pancreas while maintaining the regenerative capacities of the pancreas. The pancreas does not have a high basal rate of cell proliferation compared to other digestive epithelia such as colon or intestine. Recent single-cell analysis uncovered that this increase of proliferating acinar cells comes from heterogenous exocrine cell subsets during pancreatic regeneration (*41*). PI3Kα genetic and pharmacological inactivation was previously demonstrated by us and others to prevent oncogenic transformation (*22*, *23*). We demonstrate here that PI3K inactivation induces an increase of the pool of proliferating cells, while Kong et al showed that proliferating acinar cells are refractory to oncogenic Kras-transformation (*34*). Understanding this balance is critical to prevent cancer formation in an inflammatory context.

Strobel et al. suggested early on that the absence of expression of the HPAP reporter in exocrine lineage after pancreatic injury was indicative that acinar-to-ductal or acinar-to-centro-acinar transdifferentiation could occur, but that this process did not contribute significantly to exocrine regeneration (*42*). While ADM participates in the intrinsic defence mechanism towards pancreatic injury by reducing intra-tissular or systemic leakage of activated digestive enzymes, we found that it also contributes to fibrosis accumulation. As stromal architecture directs early dissemination in pancreatic ductal adenocarcinoma (*43*), any strategy that prevent fibrosis accumulation in the tissue could contribute to protect from the development of PDAC. In our previous work, we showed that GDC-0326 treatment after induction of PanINs by mutant KRAS reverts PanIN formation and renormalizes established fibrosis accumulation (*24*), further highlighting the importance of this interception strategy.

There is an extensive body of literature showing that increased PHGDH catalytic activity is important for (cancer cell) proliferation (*44–46*). Our findings add a new level of understanding to the biology of PHGDH by showing that low level PHGDH protein expression is associated with cells undergoing pancreatic plasticity. This finding does not contradict previous data but echoes with the recent data on metastatic breast cancer (*47*): PHGDH-low cancer cells in the primary tumour are in a slow cycling state, but have increased metastatic behaviour, usually linked to plastic properties such as epithelial-to-mesenchymal transdifferentiation potential. Overall, we highlight the importance of differential regulation of PHGDH protein level in cell plasticity.

Cancer interception is the active way of combating cancer and carcinogenesis at earlier stages. Acinar cells, albeit to a lesser extent than ductal cells (*48*), are intrinsically refractory to cancerogenesis; expression of oncogenic KRAS is required for cell transformation, but is not sufficient to drive cancerogenesis (*4*, *49*). Small molecule inhibitors of all PI3K and selective of PI3Kα are currently in various phases of clinical trials (*50*) and could be used to prevent cell plasticity while maintaining regenerative potential of pancreas. We hope that, with the rapidly developing techniques allowing detection of circulating premalignant cells and fragmented DNA (*51*) that allow to detect early disease also in preclinical models (*24*), we will be able to propose strategies to prevent pancreatic cancer in population at risk. Pharmacological inhibition of pro-cancer pathways such as those driven by PI3Kα could then be tested as a preventive strategy in patients at risk for pancreatic cancer development.

## Material & Methods

### Nutrition of Rats

Five-week-old male Fischer 344 rats were purchased from Charles River (France). Animal experiment was authorized by the French Ministry for Higher Education, Research and Innovation (MESRI) in accordance with the local ethic committee evaluation (APAFiS #16139-2018071617502938v3). Rats were randomly assigned to their respective groups and cages randomly placed in the animal facility. They were allowed to one-week acclimatization with Control diet before being i.p. injected with azoxymethane (Sigma-Aldrich) (20mg/kg) in buffered saline solution, in order to initiate colorectal cancer, as those rats were also used for a colorectal cancer study (*52*). Experimental diets started one week after azoxymethane injection, for 100 days. Rats had unlimited access to tap water and diet, given every day just before rat active period to limit food oxidation. All diets were based on modified low calcium AIN-76 diets with two powder composition: one for rat acclimatization and Control diet group with 32% protein as casein, and one, with no protein, to complement meat in the Meat diet group. These powders were prepared by SAAJ (INRAE, Jouy-en-Josas, France). All diets contained 5% safflower oil (MP Biomedicals 102888). Meat diet included pieces of cooked red meat (87,6 % of meat) and ham (12.4 % of meat) (beef sirloin was purchased from Picard, France and cooked ham from “Votrecharcutier.fr”, France). Both diets were balanced for protein and lipid contents (the control diet contained 10.16% of lard for this purpose). After 100 days of experimental diet, pancreas were collected.

### Transcriptomics gene signatures

PI3Kα gene signature was designed as the intersection of genes up- and down-regulated by shRNA against PIK3CA in a human PIK3CA mutated breast cancer cell line (*53*) and PI3ka_human_LINCS_CMAP and PI3Ka_human_mTOR_CMAP LINCS gene signatures (*54*), as described in (*24*).

### Reagents

Reagents were purchased as follows: BYL-719 (MedChem Express), CBR-5884 (MedChem Express), NCT-503 (MedChem Express), Caerulein from Bachem (ref. 4030451), Dexamethasone (Sigma-Aldrich, D1756), EGF (R&D Systems, 236-EG-200), Tamoxifen (MedChem Express). GDC-0326 was a gift from Genentech (*in vitro* IC50 in nM: p110α: 0.2; β: 133; δ: 20; γ: 51; C2β: 261; C2α>10µM; vps34: 2840; K_i_ mTOR in nM: 4300)(*55*). 1938 was provided by Cancer Tools.

### Genetically engineered mouse models

The LSL-Kras^G12D^ (from D Tuveson, Mouse Models of Human Cancers Consortium repository (NCI-Frederick, USA), Pdx1-cre (from DA Melton, Harvard University, Cambridge, MA, USA)(*56*), p110α^lox^ (from B. Vanhaesebroeck, UCL, London), Elastase CreERT2 mice (Elas-CreERT2) were generated by DA Stoffers (Philadelphia, PA, USA) and were kindly provided by Pr P Jacquemin (UCL, Belgium). Strains were interbred on mixed background (CD1/C57Bl6) to obtain compound mutant Pdx1-Cre;LSL-Kras^G12D^ (named KC), pdx1-Cre;p110α^lox/lox^ (Pdx1-Cre;p110α^lox/lox^ were named α^in^C; pdx1-Cre;LSL-Kras^G12D^;p110α^lox/lox^ were named Kα^in^C) or Elas-CreERT2; p110α^WT/WT^ (CT mice), Elas-CreERT2; p110α^lox/lox^ (α^in^E mice). For experiment with Pdx1-Cre, littermates not expressing Cre as well as Pdx1-Cre and p110α^lox/lox^ mice of the same age were used as control; for experiments with Elastase CREERT2, CT mice injected with tamoxifen are controls. Seven week-old α^in^E and CT mice were treated with tamoxifen (Sigma-Aldrich) dissolved in corn oil containing 10% ethanol at a concentration of 20 mg/ml. Mice received one intra-peritoneal (i.p.) injections (1 mg/15 g of body weight) on three consecutive days (*57*). All procedures and animal housing are conformed to the regulatory standards and were approved by the Ethical committee according to European legislation translated to French Law as Décret 2013-118 1st of February 2013 (APAFIS 3601-2015121622062840; APAFIS#32411-2021070616504074).

### Statistics

Experimental data provided at least three biological replicates. Data were analysed by 2-tailed, unpaired Student’s t-test or ANOVA test using a multiple statistics Graph Pad Prism software package. A difference was considered significant when p value was lower than 0.05: * P < 0.05, ** P < 0.01, *** P < 0.001. Non-significant (ns) if P > 0.05.

All authors had access to the study data and had reviewed and approved the final manuscript.

## Acknowledgments

We thank all members of SigDYN, past and present, for their technical support, sample banks, common tools, scientific and protocol discussions, Arnaud Besson for IHC p27 antibody and stainings, Anexplo team work, Christèle Segura (I2MC, Inserm 1048, Toulouse, France) for part of histology sample preparation, EZOP for Inrae animal facility, Imagin (FX Fresnois) for image scanning, the CRCT core technology platform, Lorie Friedman (Genentech) for their kind gift, Pierre Cordelier for access to its microscope, Anne Couvelard for explaining the scoring method of pancreatitis and taking time to read the slides with us. JGG, MD were members of COST action EU-Pancreas BM1204. Rats from red-meat enriched diet and contraol diet were from the study funded by JPI HDHL « Meatic » https://www.healthydietforhealthylife.eu/76-hdhl-intimic-cofunded-call/393-meatic; Faecal Microbiome as determinant of the effect of diet on colorectal cancer risk: comparison of meat based versus pesco-vegetarian diets; pancreas were dissected at the same time for this study. JGG’s laboratory belongs to Toucan, Laboratoire d’Excellence, ANR, an integrated research program on Signal-targeted Drug Resistance. JGG’s laboratory for this topic was/is funded by Europe EU-ERG FP7 (270696 PaCa/PI3K), ARC (PJA2021060003932, PJA20171206596, PJA20141201744; salary for RB), Toucan ANR Laboratory of Excellence, MSCA-ITN/ETN PhD-PI3K (Project ID: 675392, salary for SA), Fondation de France (salary for BT), ANR-JCJC (RADIANCE-salary for BT), GSO, Ligue Nationale Contre le Cancer, Arc (salary for CC). SA disseminated their research to high school students, as part of their commitment in MSCA-ITN funding. DB and AVV were funded by Erasmus + program. F Pierre, F Guéraud have some research projects from their academic research teams that have been co-financed by the processed meat sector. The other authors declare no conflict of interest.

## Supplementary Material and methods

### Bioinformatics analysis

Amongst the publicly available micro-array repositories, we selected transcriptional profiling datasets of normal pancreas, chronic pancreatitis and pancreatic cancer tissues, including 8 normal, 9 pancreatitis (alcoholic or autoimmune), 5 stroma of chronic pancreatitis (undocumented aetiology) and 7 pancreatic adenocarcinomas. Published data on human samples were retrieved from public databases (E6MEXP-804(58), E-MEXP-1121(60), E-TABM-145 (59)) from compatible platforms, normalized using RMA method (R 3.2.3, bioconductor version 3.2), collapsed (collapse microarray), filtered (SD>0.25), and statistically tested using an ANOVA test corrected with Benjamini & Hochberg method (BH). Published murine mRNA expression during experimental pancreatitis (GSE65146) was analysed by TTCA package of R (kindly corrected by its conceptor) and normalised with SCAN(61). For each sample, individual scoring for hallmarks or Reactome (actualised list of genes downloaded from MSigDB version [software.broadinstitute. org/gsea/msigdb] and Reactome [www.reactome.org]) was performed using Autocompare_SES software (available at https://sites.google.com/site/fredsoftwares/products/autocompare_ses) using the “greater” (indicating an enriched gene set) Wilcoxon tests with frequency-corrected null hypotheses (62), followed by values in each compared group using an ANOVA test. Hierarchical sample clustering was performed using the PI3Kα activation signature.

### Genotyping

Genotyping was performed as described in (22, 24): the genetically modified allele is called p110α^lox^ or p110α^ΔDFG^ (here named Rec) after Cre recombination. DFG is a conserved motif in the activation loop of the p110α kinase domain critical for its catalytic activity. The gene targeting strategy used is different from a traditional conditional knock-out strategy. Instead, it is deleting two exons in the catalytic domain of Pik3ca in the 3’ part of the gene, allowing expression of a truncated inactive p110α after recombination. Primers corresponds to post-cre primers used to verify the presence of recombined allele ΔDFG: ma9: ACACACTGCATCAATGGC; ma5: GCTGCCGAATTGCTAGGTAAGC; annealing temperature: 65°C; recombined ΔDFG: 544 bp, wild type (+) or unrecombined lox: >10kbp amplicon. Genotyping of tail or pancreas samples was performed after DNA extraction with Sigma kit (XNAT-100RXN). Genotyping primers for Elastase CreERT2 are: ElasCreERT2 ElasF-CTCTGCTAACCATGTTCATGCCT ElasR-ACGCTAGAGCCTGTTTTG.

### Preparation of drug treatments

For in vitro experiments, GDC-0326 powder is resuspended at 10mM in DMSO then diluted as indicated. BYL-719 power is resuspended at 10mM in DMSO then diluted as indicated. CBR-5884 and NCT-503 powders are resuspended at 10mM in DMSO, sonicated at 4°C for 10min (10 cycles 30sec sonication + 30 sec stop) then diluted in medium as indicated.

For in vivo experiments, the powder is dissolved in 0.5% methylcellulose containing 0.1% Tween 80 or 0.2% (for 1938 only) (Sigma) using vortex and rotate at 4°C for 15 minutes. The solution is then sonicated at 4°C for 10min (10 cycles 30sec sonication + 30 sec stop). Solution is transferred into a wheaton vial. The solution is mixed well before administration. Dosing volume is 4mL/kg for PO dosing.

### Mouse housing

The animals were housed under the following conditions. The animal facility has controlled access. Housing rooms are maintained at a constant temperature of between 20 and 22°C. A 12/12h light cycle is automated. Animals are housed in GM500 cages on ventilated Tecniplast Green Line racks. The animals are provided with an enriched environment consisting of absorbent paper and an igloo for nesting. The following procedures were used: use of the tunnel to refine the grip (or cupped hand…); animal training and habituation methods before experimenting; environmental enrichment provided to the animals; daily visits by trained and competent staff; limiting points and scoring grid to ensure optimal and rapid decision. General discontinuation criteria are: weight loss of up to 20%, severe abnormal behaviour, cessation of feeding, a drop in body temperature, or mutism lasting more than 24 hours, which never occurred. In such cases, the animal would have been immediately euthanized. Specific pain management: final blood sampling procedures, anesthesia, followed by terminal anesthesia. The ARRIVE reporting guidelines were used to describe the preclinical models.

### Repetitive caerulein-induced pancreatic injury & PI3K inhibitor/activator in vivo dosage

Total number of mice for each genotype was produced. After genotyping, when indicated, tamoxifen dosage was performed blindly in all the cages, where animal are grouped by litter or concomitant litter. Next, mice of each genotypes were randomly assigned to their respective treatment groups and grouped by cages of treatment (separating males and females without any animal being left alone). Cages were randomly placed in the animal facility. Mice were weighed one day prior experiment to allow calculation of drug and caerulein dosage; the order of the cage treatment was random. Pancreatic injury was induced on young mice (8-12 weeks) by series of six hourly intra-peritoneal injections of caerulein (75µg/kg of body weight) or saline that was repeated every 2 days in the presence or absence of GDC-0326 (10mg/kg). Animals were euthanized at 7 days, and recombination was verified before analysis as described in (32, 38). GDC-0326 (10mg/kg) was administrated by gavage every morning. Amylase measurement was performed using Phadebas kit (Phadebas Magle, used as suggested) in plasma samples at early time points. As a read-out of efficient pancreatic PI3Kα inhibition, glycemia was determined from tail blood using a glucose monitor (Accu-Chek Performa). 2.5 weeks after intraperitoneal tamoxifen injection two rounds of caerulein 8-hourly intraperitoneal injections was performed at day 1 and day 3 (the animals fasted the night before caerulein injection). In the same time mice are treated by gavage twice a day for 3 days with vehicle or 1938 at 100mg/kg. No mice was excluded in those experiments. A priori sample size was performed using previous experiments for IHC characterisation of differences in cell proliferation rate or immune reaction (primary outcomes), as the estimated number of subjects needed in each group in order to demonstrate a statistically significant difference at “p” values ranging from 0.05 - 0.01 and at 95% levels of “power” was 4-6 (24). One main limitation of the proposed caerulein model is that it mimics the early steps of inflammation; the setup of injection did not allow to reach the full chronic pancreatitis stage observed in Humans.

### Histology and Immunostaining

Immunostainings were conducted using standard methods on formalin-fixed, paraffin-embedded tissues, including both mice and human pancreas. Antigen retrieval and antibody dilution was carried out as described in Supplementary Table 2. Picrosirius Red was from from Sigma, Direct Red 80 (used as suggested and analysed under polarized light.

Specific following settings were used for quantification of Ki67 (5 slides at 10x / mouse; Image J: Threshold Color, 8bit, Threshold, Watershed, Compting particles (0-infinity for pixel number, circularity: 0.1-1). The other qualifications were done manually in a blind fashion with slides ranged per intensity prior starting of the quantification. The persons that analysed the slides were different from the persons that performed the experimental setting.

### In vitro culture of acinar cells and treatments

AR42J cells from ATCC are a rat pancreatic acinar cell line. They were cultured in Advance Dulbecco’s Modified Eagle’s Medium F12 containing 10% FBS (Eurobio), 1% L-glutamine (Sigma-Aldrich) 1% Peniciline-streptomycin (Sigma-Aldrich). Acinar-to-ductal transdifferentiation was induced with 1 µM Dexamethasone and 10 ng/ml EGF (for 8 days). The culture medium containing diluted agents and inhibitors was changed every 48 h.

### RT-qPCR analyses

Cells were harvested on ice, washed twice with cold PBS, collected, and frozen at -80°C. RNA of cell pellets was isolated according TRIzol protocol (Life Technologies). RT-qPCR reactions were carried out using RevertAid H Minus Reverse Transcriptase (Thermo Fisher) and SsoFast EvaGreen Supermix (Bio-Rad) according to the manufacturers’ instructions. The list of primers is in Suppl Table 2.

### Western blot analysis

Cells were harvested on ice, washed twice with cold PBS, collected, and frozen at -80°C. Dry pellets of cells were lysed in 50 mM Tris-HCl, pH 7.5, 150 mM NaCl, 1 mM EDTA, 1% Triton-X100 (Sigma-Aldrich) supplemented with protease and phosphatase inhibitors (sodium orthovanadate (Sigma-Aldrich), 1 mM DTT, 2 mM NaF (Sigma-Aldrich) and cOMPLETE Mini Protease Inhibitor Cocktail (Roche). Derivatization of carbonyl groups in protein lysates was performed according to standard protocol (Ab178020). Protein concentration was measured using BCA Protein Assay kit (Interchim), and equal amounts of proteins were subjected to SDS-PAGE and transferred onto nitrocellulose membrane (BioTraceNT; Pall Corp). Membranes were washed in Tris Buffered Saline supplemented with 0.1% Tween-20 (TBS-T) then saturated in TBS-T with 5% non-fat dry milk, incubated overnight with primary antibodies in TBS-T with 5% BSA, washed and revealed according to Cell Signaling Technology protocol. Western blotting was conducted using standard methods with antibodies as described in the Supplementary Table 2.

### Biochemical analysis of PHGDH activity

Protein extraction was performed on cells using cold PHGDH assay buffer. Protein concentration was determined using BCA assay. Protein precipitation was performed using protein extract and saturated ammonium sulfate solution. PHGDH activity is determined using PHGDH Assay kit from Abcam ab273328, following the instructions, with PHGDH developer solution and PHGDH substrate. The luminescence is followed using CLARIOstar spectrophotometer at 37°C at 450nm.

### Real-time metabolic analysis and XF Cell Mito Stress Test (OXPHOS experiment)

Measurements were performed using the Seahorse Bioscience XFe24 Extracellular Flux Analyzer (Agilent). This device allows the measurement of the cellular oxygen consumption rate (OCR in pmoles/min) and of the extracellular acidification rate (ECAR in mpH/min), for mitochondrial respiration (OXPHOS) and glycolysis, respectively. Forty-eight hours before the assay, cells at exponential growth were seeded into Seahorse 24-well plates and cultured at 37°C with 5% CO_2_. The number of seeded cells was optimized to ensure 70%–80% confluence the day of analysis. Twenty-four hour after seeding, cells are treated with the indicated combination of inhibitors and incubated at 37°C with 5% CO_2_.

OCR was measured using the Seahorse XF Cell Mito Stress Test Kit. Culture medium was replaced with OXPHOS assay medium (Seahorse XF base medium, 2 mM glutamine, 1 mM sodium pyruvate and 10 mM glucose, pH 7.4) and the plate was pre-incubated for 1h at 37°C in an incubator without CO_2_. OCR was measured under basal conditions, and then after sequential injections of different reagents: 1 µM oligomycin (respiratory Complex V inhibitor that allows to calculate ATP production by mitochondrion), carbonyl cyanide-p-trifluoromethox-yphenyl-hydrazon (FCCP; an uncoupling agent allowing determination of the maximal respiration and the spare capacity; 1µM), and finally 0.5 µM rotenone + 0.5 µM antimycin A (Complex I and III inhibitors, respectively) to stop mitochondrial respiration enabling the calculation of the background (i.e., non-mitochondrial respiration driven by processes outside the mitochondria). Levels of OCR were normalized to 25,000 seeded cells, which we found as the most accurate way to normalize when comparing different cell types.

### siRNA transfection

Cells were transfected with Lipofectamine RNAiMAX (ThermoFisher Scientific) with SMARTpool ON-TARGETplus rat siRNA (Dharmacon) targeting: Phgdh according to the manufacturer’s protocols as used in(24). ON-TARGETplus Non-targeting control siRNAs (Dharmacon) were used as controls. Forty-eight hours after transfection, the cell protein contents were extracted and analysed by western blot to determine the efficacy of siRNA transfection using anti-PHGDH antibody.

### Cell processes analysis

Cell proliferation/morphological changes are estimated using the confluence followed by phase microscopy using Incucyte. Cells are incubated for seventy-two hours in 96 well plate. Two pictures by well are taken using 10x objective. Using Incucyte software the cell surface is measure on the picture, a mean of each replicate is used to compare the different condition at the different time points.

### Acinar cell isolation

Mice were anesthetized with isofluorane and sacrificed by cervical dislocation. Pancreas were quickly resected, rinsed and 100 mg of the organ were placed in 5 ml of cold HBSS (Hank’s balanced salt solution) containing 0.01% (w/v) Soybean Trypsin Inhibitor (STI, Sigma-Aldrich) and 1000U of collagenase II (Thermofisher). Digestion was performed at 37°C during 20 minutes and mechanical dissociation every 5 minutes through 10 back-and-forth passages of the pancreatic tissue into sterile serological pipettes of decreasing size (25, 10 and 5 ml). Following multiple washes with HBSS supplemented with 5% FCS, 0.01% STI and 10 mM Hepes (pH 7.5), digested pancreatic tissue was pelleted by 400 rpm centrifugation, resuspended in DMEM culture medium containing 4.5 g/l glucose, 10% fetal calf serum (FCS) (Eurobio), 10% penicillin-streptomycin mixture, 0.01% STI and filtered through 100 µm-Nylon mesh.

### EdU assay

Isolated acinar cells were placed in Ibidi 8 well plate coated with gelatine at 37% and 5% CO2. After 48h plating, cell are treated for 24h using the indicated inhibitor combination. To determine the inhibitor effects on cell proliferation, cells are incubated with 10µM EdU solution in the last 2 hours at 37°C with 5% CO_2_. Cells are fixed using PFA for 15min and permeabilised using a solution of PBS containing 0.5% Triton-100x for 20 min. The EdU integration is revealed using the indicated solution. DAPI coloration was performed for 1min to quantify all nuclei. Edu incorporation is observed using Operetta immunofluorescence microscope. EdU positive nuclei were quantified using Harmony software.

Suppl. Fig. 1 **Supplementary data related to Figure 4**. PHGDH inhibitors were added in vitro in extracts of acinar AR4-2J cells. **b.** Genotyping of cre-recombined pik3ca allele in Cre negative or Pdx1-Cre;p110α^lox/lox^ (α^in^C) pancreata. Representative results in 4 different mice are shown; the amplicon corresponds to the recombined allele (Rec) which is absent in WT pancreas. **c.** α-Amylase measurements in 10µL plasma diluted 1/20 times was performed at indicated times. α-Amylase activity in arbitrary unit is shown and represented as mean ± SEM. T-Test: **=P<0.01. **d-e.** IHC staining as indicated in WT or α^in^C pancreas subjected to the repeated injury chronic pancreatitis protocol (Caerulein). Scale: 20µm. Representative pictures are shown.

Suppl. Fig. 2 **Supplementary data related to Figure 5. a.** IHC staining as indicated in WT or α^in^C pancreas subjected to the repeated injury chronic pancreatitis protocol (Caerulein). Scale: 20µm. Representative pictures are shown. b. H&E WT or α^in^C pancreas subjected to the repeated injury chronic pancreatitis protocol (Caerulein). Scale: 50µm.

**Supplementary Table 1** Statistical analysis of Hallmarks gene signatures and PI3Kα-regulated gene expression in normal, CP (from alcoholic or auto-immune origin), PDAC or microdissected stroma of CP patients. Assembled mean values of alcoholic CP, autoimmune CP, CP stroma, normal or PDAC pancreas are shown; p-value and BH-corrected p-value are shown for PDAC vs. normal, or all vs. normal statistical tests. Lists of genes as indicated.

Supplementary Table 2 **List of tools and protocols.**

## Notes

### Competing Interest Statement

F Pierre, F Gueraud have some research projects from their academic research teams that have been co-financed by the processed meat sector. The other authors declare no conflict of interest.

### Summary of Updates

- further quantifications were added to our meat enriched diet protocol (fig. 1) - further in vitro demonstration for the role fo PHGDH in determining the signalling output of PI3K in exocrine pancreas was added (fig. 3, fig. 4) - further analysis of all preclinical models of pancreatitis was added (fig. 5) - with those new data, title, abstract and discussion were also updated

