## Supplementary figures and images for "Meat enriched-diet and inflammation promote PI3Kα-dependent pancreatic cell plasticity that limit tissue regeneration"

### FIGS1

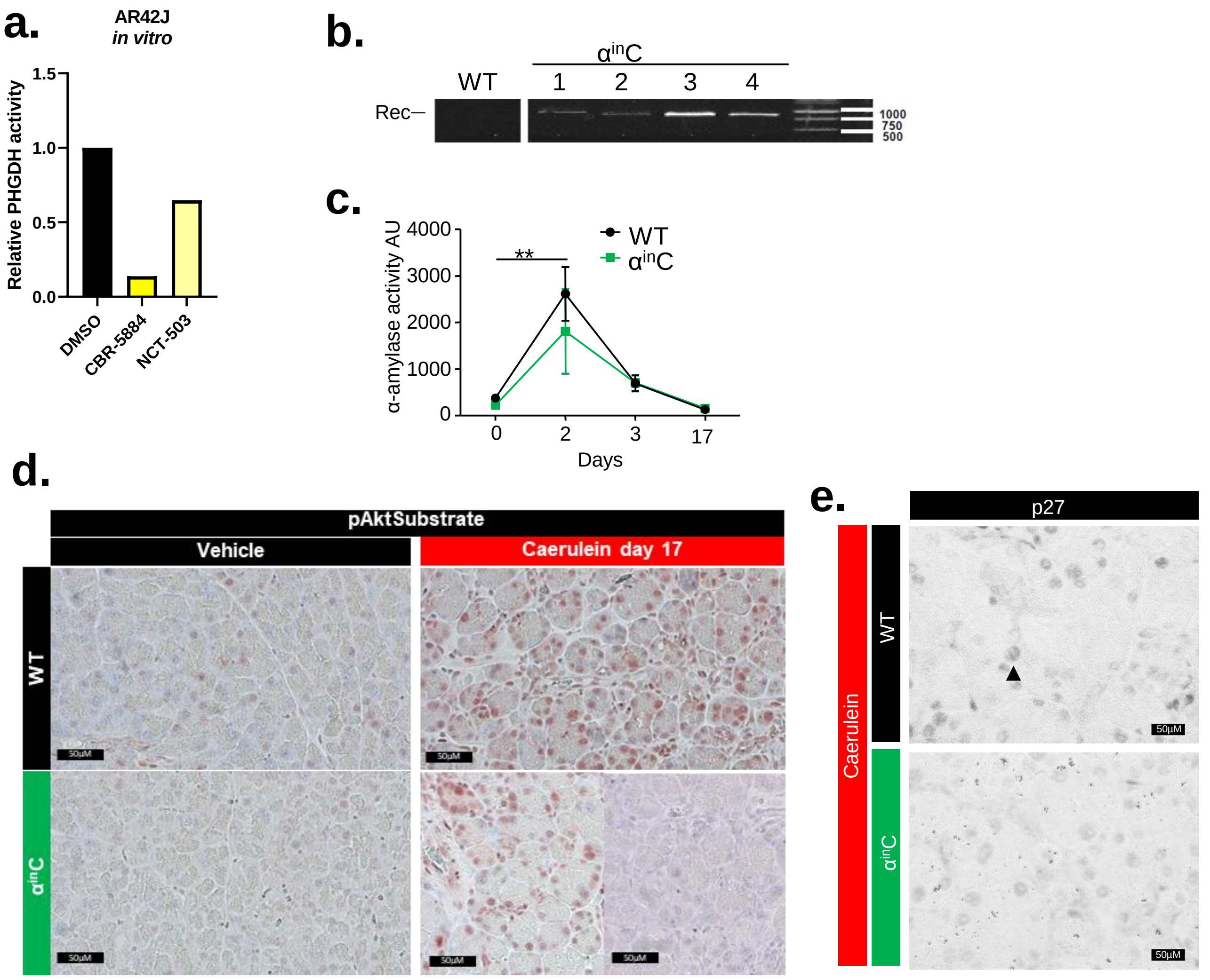


#### Figure Suppl 1

### FIGS2

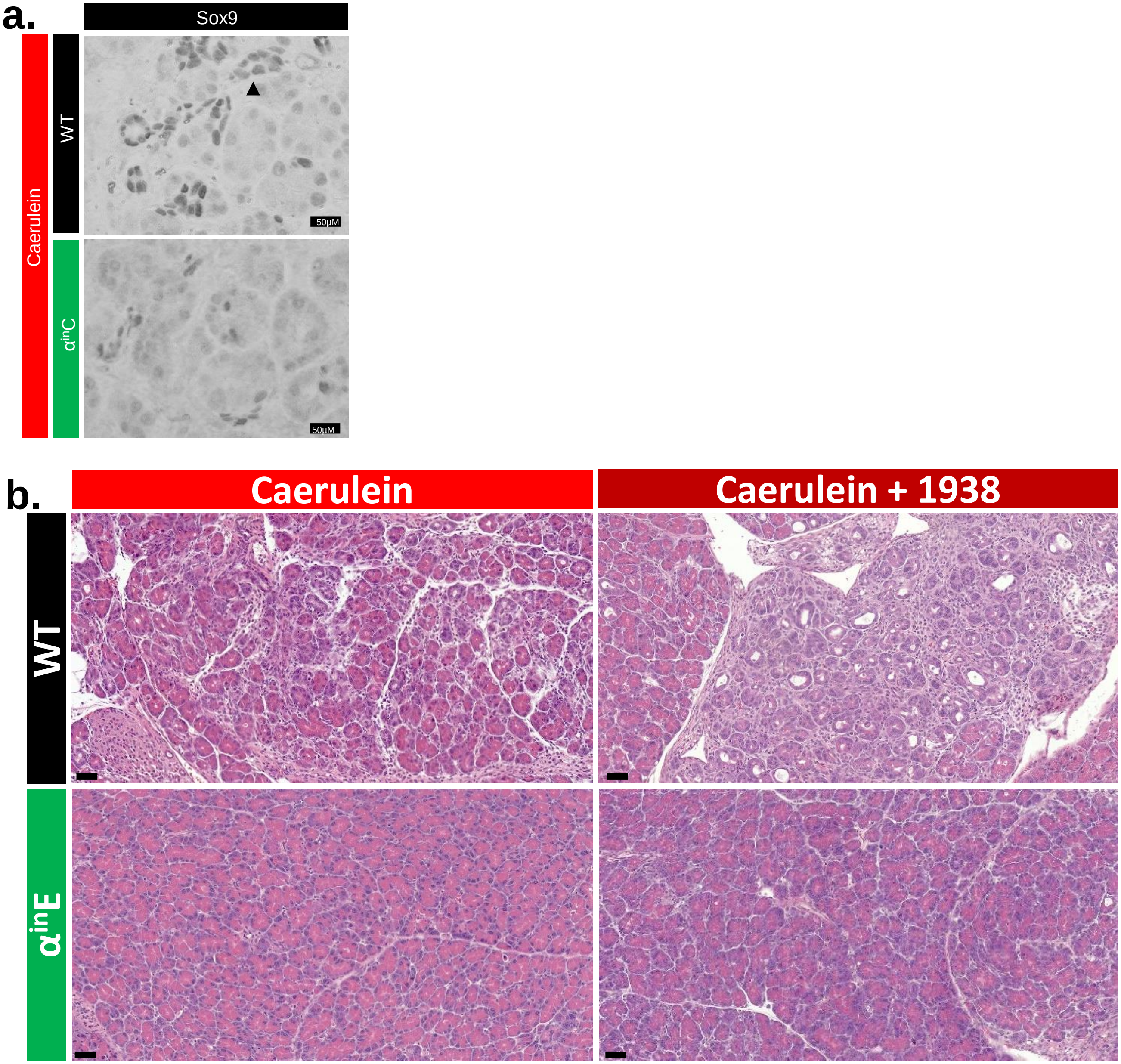


#### Figure Suppl 2
